# Syngap1 and the development of murine neocortical progenitor cells

**DOI:** 10.1101/2024.12.18.629233

**Authors:** Soraia Barão, Ingie Hong, Ulrich Müller, Richard L. Huganir

**Author notes:** Corresponding authors: Richard L. Huganir, Ulrich Müller and Ingie Hong.; and.

## Abstract

SYNGAP1 is a major regulator of synaptic plasticity through its interaction with synaptic scaffold proteins and modulation of Ras and Rap GTPase signaling pathways. *SYNGAP1* mutations in humans are often associated with intellectual disability, epilepsy, and autism spectrum disorder. *Syngap1* heterozygous loss-of-function results in impaired LTP, premature maturation of dendritic spines, learning disabilities and seizures in mice. More recently, SYNGAP1 was shown to influence cortical neurogenesis and the proliferation of progenitors in human organoids. Here, we show that the absence or haploinsufficiency of *Syngap1* does not influence the properties of neocortical progenitors and their cellular output in mice. This discrepancy highlights potential species-specific or methodological differences and raises important questions about the broader applicability of SYNGAP1’s role in neurogenesis.

## Introduction

Synaptic Ras GTPase-activating protein 1 (SYNGAP1) is a critical postsynaptic protein predominantly expressed in the brain, where it plays a crucial role in synaptic plasticity, learning, and memory^1,2^. Since its discovery in the late 1990s^3,4^, SYNGAP1 has been identified as a major regulator of synaptic strength through its interaction with synaptic scaffold proteins and modulation of Ras and Rap GTPase signaling pathways. It is one of the most abundant proteins in the postsynaptic density (PSD)^5–7^, where it interacts with PSD-95 and other critical scaffolding proteins, positioning it as a pivotal component in regulating excitatory synapses. SYNGAP1’s GAP activity modulates Ras-ERK and PI3K-Akt signaling pathways, which can regulate synaptic plasticity mechanisms, including long-term potentiation (LTP)^8^.

Research in knockout mice showed that *Syngap1* heterozygous loss-of-function (LoF) results in strongly impaired LTP, premature maturation of dendritic spines, and learning disabilities^9–11^. Later studies identified spontaneous seizure activity and seizure susceptibility in these mice^12,13^. Homozygous knockout mice died within days of birth^9–11^ co-incident with the time frame when the expression of *Syngap1* dramatically increases in the brain^14,15^. These structural and functional abnormalities underline the importance of SYNGAP1 in postnatal neurodevelopment and synapse regulation, supporting its role as a critical molecular hub in synaptic function. Following the identification of SYNGAP1’s role in synaptic plasticity, heterozygous LoF mutations in human *SYNGAP1* were linked to a highly penetrant and severe neurodevelopmental disorder (SYNGAP1 syndrome; MRD5; SYNGAP1-related intellectual disability; SYNGAP1 encephalopathy)^16–18^. Patients with *SYNGAP1* mutations often present with intellectual disability, epilepsy, and autism spectrum disorder. With more than 1,400 genetically diagnosed patients worldwide and an estimated prevalence at around 6 out of 100,000 individuals^19^, *SYNGAP1* has become a key target for understanding the molecular mechanisms that contribute to neurodevelopmental diseases and cognitive impairment.

The majority of these studies have focused on developmental stages after neuronal migration is complete and when excitatory synapses have started to emerge. More recently, studies have begun exploring the potential involvement of SYNGAP1 in earlier stages of neurodevelopment, including cortical development^20,21^. Willsey et al.^20^ showed that *SYNGAP1* depletion leads to an increased ratio of neural progenitor cells to neurons and a reduction of ventricle and telencephalon size in *Xenopus*. Based on work with human cortical organoids, Birtele et al.^21^ concluded that *SYNGAP1* haploinsufficiency accelerated neuronal maturation which decreased the ratio of progenitor cells to neurons and resulted in a reduced ventricular area in this system. These findings suggest that *SYNGAP1* regulates developmental processes during brain development at much earlier stages than previously thought.

In this study, we sought to determine the extent to which these results extend to cortical neurogenesis in mice. Using histological analysis of genetically modified mice carrying heterozygous and homozygous knock-out alleles for *Syngap1 (Syngap1KO*), we did not observe significant alterations in the properties of cortical progenitors and the generation of neocortical cell layers. We thus conclude that *Syngap1* is not essential for neocortical development in mice. Our findings highlight potential species-specific or methodological differences and raise important questions about the broader applicability of *SYNGAP1*’s role in neurogenesis.

## Results

### *Syngap1* levels do not affect the number of neocortical progenitor cells in mice

Recent studies have proposed that *SYNGAP1* plays a role in cortical neurogenesis by affecting the proliferation and differentiation of progenitor cells in human organoids^21^. Cortical progenitors have been broadly divided into two classes named apical progenitors (APs = PAX6-positive cells) that undergo mitosis in the ventricular zone (VZ), and intermediate progenitors (IPs = TBR2-positive cells) that undergo mitosis in the subventricular zone (SVZ)^22–27^. Although *SYNGAP1* is mainly expressed in neurons at embryonic and postnatal ages^14,28,29^, we show by RNAscope *in situ* hybridization that some *Syngap1* mRNA is expressed in *Pax6*-positive murine neocortical progenitor cells already at embryonic days 14.5 and 18.5 (E14.5 and E18.5; Extended Data Fig. 1a; Fig. 1a). However, quantitative analysis of histological sections revealed that at E18.5 the number of PAX6^+^ APs and TBR2^+^ IPs was not altered in the somatosensory cortex of heterozygous and homozygous *Syngap1KO* mice^10^ (Fig. 1b-c; rostral➔caudal axis – Extended Data Fig. 2a). The number of neocortical progenitor cells was also normal in mice carrying the *GAP-AL* mutation which eliminates the GAP activity of Syngap1 (*Syngap1GAP\**; Extended Data Fig. 2b)^8^. In addition, the cortical thickness remains unaltered in both mice at E18.5 (Fig. 1d; Extended Data Fig. 2c).

**Figure 1.**
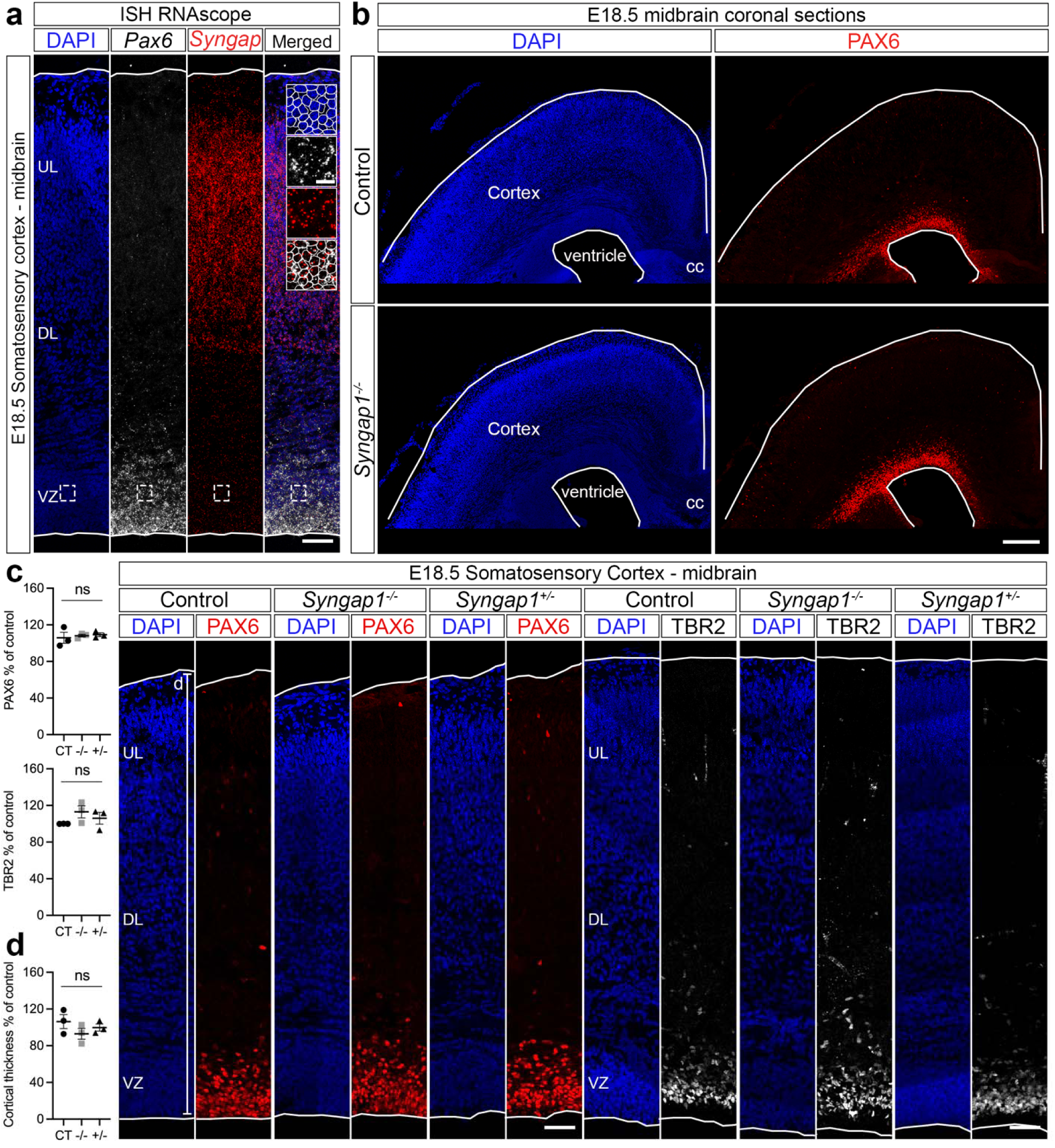
*Syngap1* levels do not affect the number of progenitor cells in mice. **a,** RNAscope for *Syngap1* (red) and *Pax6* (grey) in the somatosensory cortex of wild-type mice at E18.5. Low and top lines represent the limits of the ventricular zone (VZ) and cortical plate (CP), respectively. Boxed area at higher magnification on the right. Lines circulating the cells show co-expression *Syngap1* and *Pax6* in the VZ. **b,** Overview of midbrain coronal sections of control and homozygous *Syngap1* knockout (*Syngap1KO*) mice analyzed by PAX6 immunolabeling at E18.5. Lines represent the limits of the ventricle and CP. **c,** Control (CT), heterozygous (+/-) and homozygous (-/-) *Syngap1KO* brains analyzed by PAX6 and TBR2 immunolabeling in the somatosensory cortex at E18.5 (n=3 mice/genotype; two-sided unpaired t*-*test: PAX6 CT vs -/-p=0.7337; PAX6 CT vs +/- p=0.6224; TBR2 CT vs -/- p=0.1170; TBR2 CT vs +/- p=0.4029). **d,** Cortical thickness in the somatosensory cortex of Control (CT), heterozygous (+/-) and homozygous (-/-) *Syngap1KO* brains at E18.5 (n=3 mice/genotype; two-sided unpaired t*-*test: CT vs -/- p=0.2374; CT vs +/- p=0.4845). DL, Deep Layer; UL, Upper Layer; cc, Corpus Callosum; Values are mean ± SEM; ns, not significant; Scale bars: 50 µm (lower magnification), 10 µm (higher magnification), 500 µm (b, overview).

### *Syngap1* levels do not affect the properties of progenitor cells in mice

To further investigate the potential role of *SYNGAP1* in progenitor function, we injected EdU into pregnant mice at E18.5 and analyzed EdU incorporation into DNA 1h later in the somatosensory cortex of heterozygous and homozygous *Syngap1KO* mice (Fig. 2a,a’). Consistent with an unaltered number of progenitor cells, the number of EdU^+^ cells was also unaffected in the somatosensory cortex of heterozygous and homozygous *Syngap1KO* mice at E18.5 (Fig. 2a). Next, we investigated cell cycle exit by injecting EdU into pregnant mice at E17.5 and analyzing the number of EdU^+^/Ki67^-^ cells 24h later (Fig. 2b,b’). The number of EdU^+^/Ki67^-^ cells was not altered in the somatosensory cortex of heterozygous and homozygous *Syngap1KO* mice at E18.5 (Fig. 2b,b’). We conclude that the cell cycle and proliferative capacity of neocortical progenitors is normal in heterozygous and homozygous *Syngap1KO* mice.

**Figure 2.**
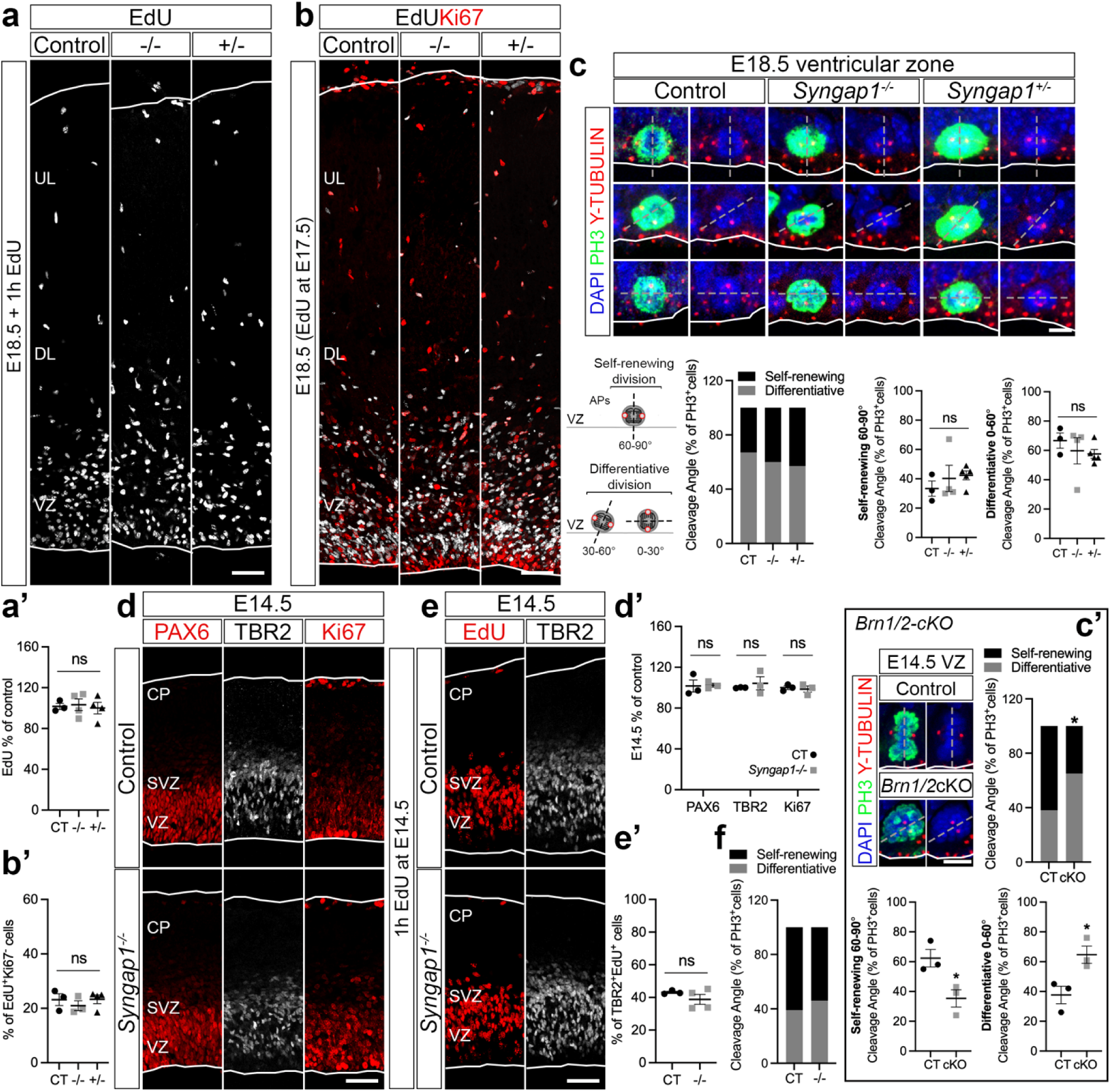
*Syngap1* levels do not affect the properties of progenitor cells in mice. **a, a’,** EdU (grey) incorporation in the somatosensory cortex of Control (CT), heterozygous (+/-) and homozygous (-/-) *Syngap1KO* mice after 1h EdU injection at E18.5 (n=3 CT, n=4 +/- and -/- mice; two-sided unpaired t*-*test: CT vs -/- p=0.8312; CT vs +/- p=0.8281). **b, b’,** EdU (grey) and Ki67 (red) immunolabeling in the somatosensory cortex of Control (CT), heterozygous (+/-) and homozygous (-/-) *Syngap1KO* mice after 24h EdU injection at E17.5 (n=3 CT and -/-, n=4 +/- mice; two-sided unpaired t*-*test: CT vs -/- p=0.4737; CT vs +/- p=0.9659). **c,** Cleavage plane analysis by PH3 (green) and γ-TUBULIN (red) immunolabeling in Control (CT), heterozygous (+/-) and homozygous (-/-) *Syngap1KO* ventricular zone (VZ) at E18.5 (n=3 CT, n=4 -/-, n=5 +/- mice; two-sided unpaired t*-*test: CT vs -/- p=0.5721; CT vs +/- p=0.1561). **c’,** Cleavage plane analysis by PH3 (green) and γ-TUBULIN (red) immunolabeling in Control (CT) and *Brn1/2- cKO* VZ at E14.5 (n=3 mice/genotype; two-sided unpaired t*-*test: CT vs cKO p=0.0308). **d, d’,** PAX6 (red), TBR2 (grey) and Ki67 (red) immunolabeling in the somatosensory cortex of Control (CT) and *Syngap1^-/-^* mice at E14.5 (n=3 mice/genotype; two-sided unpaired t*-*test: PAX6 p=0.9060; TBR2 p=0.5374; Ki67 p=0.7142). **e, e’,** TBR2 (grey) and EdU (red) immunolabeling in the somatosensory cortex of control (CT) and *Syngap1^-/-^* mice at E14.5 (n=3 CT, n=4 -/- mice; two-sided unpaired t*-*test: p=0.2898). **f,** Cleavage plane analysis in Control (CT) and homozygous (-/-) *Syngap1KO* ventricular zone (VZ) at E14.5 (n=3 CT, n=4 -/- mice; two-sided unpaired t*-*test: CT vs -/- p=0.3010). Low and top lines represent the limits of the VZ and cortical plate (CP), respectively. DL, Deep Layer; UL, Upper Layer; SVZ, Subventricular Zone. Values are mean ± SEM; ns, not significant; *p*<*0.05; Scale bars: 50 µm (lower magnification), 5 µm (c).

Birtele et al.^21^ have shown that *SYNGAP1* haploinsufficiency affects the division mode of human progenitor cells. In the VZ, depending on their cleavage plane, APs undergo either symmetric proliferative division (vertical = 60-90 degree plane) or asymmetric differentiative divisions (oblique = 30-60 degree plane and horizontal = 0-30 degree plane respectively)^30–33^. In addition, APs engage in two modes of neurogenesis termed direct and indirect neurogenesis^22–27^. During direct neurogenesis, APs divide asymmetrically to self-renew and to generate one neuron, while during indirect neurogenesis, APs divide asymmetrically to self-renew and generate an IP that then gives rise to neurons^22–26^. Although indirect neurogenesis generates neurons for all cortical layers, it is the predominant neurogenic mode that produces upper layer projection neurons (ULNs)^23,24,34–37^. Defects in the division mode of neocortical progenitors alter the identity and topography of cortical neurons and can cause several neurological and psychiatric disorders^27,38,39^. Birtele et al. described an increased proportion of progenitors undergoing differentiative division in *SYNGAP1* haploinsufficient human organoids^21^. To determine if a similar phenotype was observed in mice, we analyzed the division planes of mitotic ventricular APs (PH3-positive cells) in heterozygous and homozygous *Syngap1KO* mice at E18.5. Quantitative analysis of histological sections in the somatosensory cortex revealed a normal ratio of self-renewing and differentiative divisions in neocortical progenitors of heterozygous and homozygous *Syngap1KO* mice (Fig. 2c). As a positive control, we analyzed *Brn1* and *Brn2* conditional knockout (*Brn1/2-cKO*) mice*. Brn1* and *Brn2* have previously been shown to regulate the proliferative capacity of cortical progenitor cells and the balance between direct and indirect neurogenesis^40^. In agreement with earlier findings, the ratio of progenitors to neurons was significantly altered in *Brn1/2-cKO* mice (Fig. 2c’), demonstrating that our analysis tools are sufficiently sensitive to detect changes in the biology of neocortical progenitor cells.

Neocortical progenitors generate neurons for different cortical layers in sequential order. Deep layer neurons (DLNs) are generated prior to ULNs, which only begin to emerge in mice around E14.5. Subsequently these progenitors start to generate glia cells around E16.5^41^. To determine the extent to which *Syngap1* affects the properties of progenitor cells during earlier developmental stages at the beginning of ULN’s generation, we analyzed markers of progenitor cells (PAX6 and TBR2) and proliferation (Ki67) in the somatosensory cortex of homozygous *Syngap1KO* mice at E14.5. Immunohistochemical analysis revealed that the number of progenitor cells as well as their proliferative capacity was normal in homozygous *Syngap1KO* mice at E14.5 (Fig. 2d,d’). Additionally, as a readout of cells actively going through indirect neurogenesis, we analyzed the number of TBR2^+^ cells that have incorporated EdU 1h after injection. Levels of indirect neurogenesis were unaltered in the somatosensory cortex of homozygous *Syngap1KO* mice at E14.5 (Fig. 2e,e’). Quantitative analysis also revealed a normal ratio of self-renewing and differentiative divisions in neocortical progenitors from the somatosensory cortex of heterozygous and homozygous *Syngap1KO* mice at E14.5 (Fig. 2f; Extended Data Fig. 1b). Altogether, these results show that the neocortical properties of progenitor cells are normal in the absence of *SYNGAP1* in mice.

### *Syngap1* levels do not affect the number of neurons and glia cells generated in mice

To further determine the extent to which *Syngap1* may have essential roles during the formation of neocortical cell layers, we analyzed numbers of ULNs, DLNs and astrocytes in heterozygous and homozygous *Syngap1KO* mice at P0 and P13. Immunohistochemical analysis of CTIP2 (highly expressed in DLNs) and BRN2 (highly expressed in ULNs at P0) did not reveal any differences in the numbers of CTIP2^+^ and BRN2^+^ neurons in the somatosensory cortex of heterozygous and homozygous *Syngap1KO* mice compared to wild-type at P0 (Fig. 3a). Numbers of CTIP2^+^ and BRN2^+^ neurons were also normal in *Syngap1GAP\** mice at P0 (Extended Data Fig. 2d,d’). In addition, we observed normal numbers of astrocytes (SOX9-positive cells) in the somatosensory cortex of heterozygous and homozygous *Syngap1KO* mice at P0 (Fig. 3b,b’). Immunohistochemistry analysis of CTIP2 and CUX1 (highly expressed in ULNs at P13) revealed normal numbers of ULNs and DLNs the somatosensory cortex of heterozygous *Syngap1KO* mice at later postnatal ages (P13; Fig. 3c). Numbers of astrocytes were also unaltered at P13 in these mice (Fig. 3d; Extended Data Fig. 2e).

**Figure 3.**
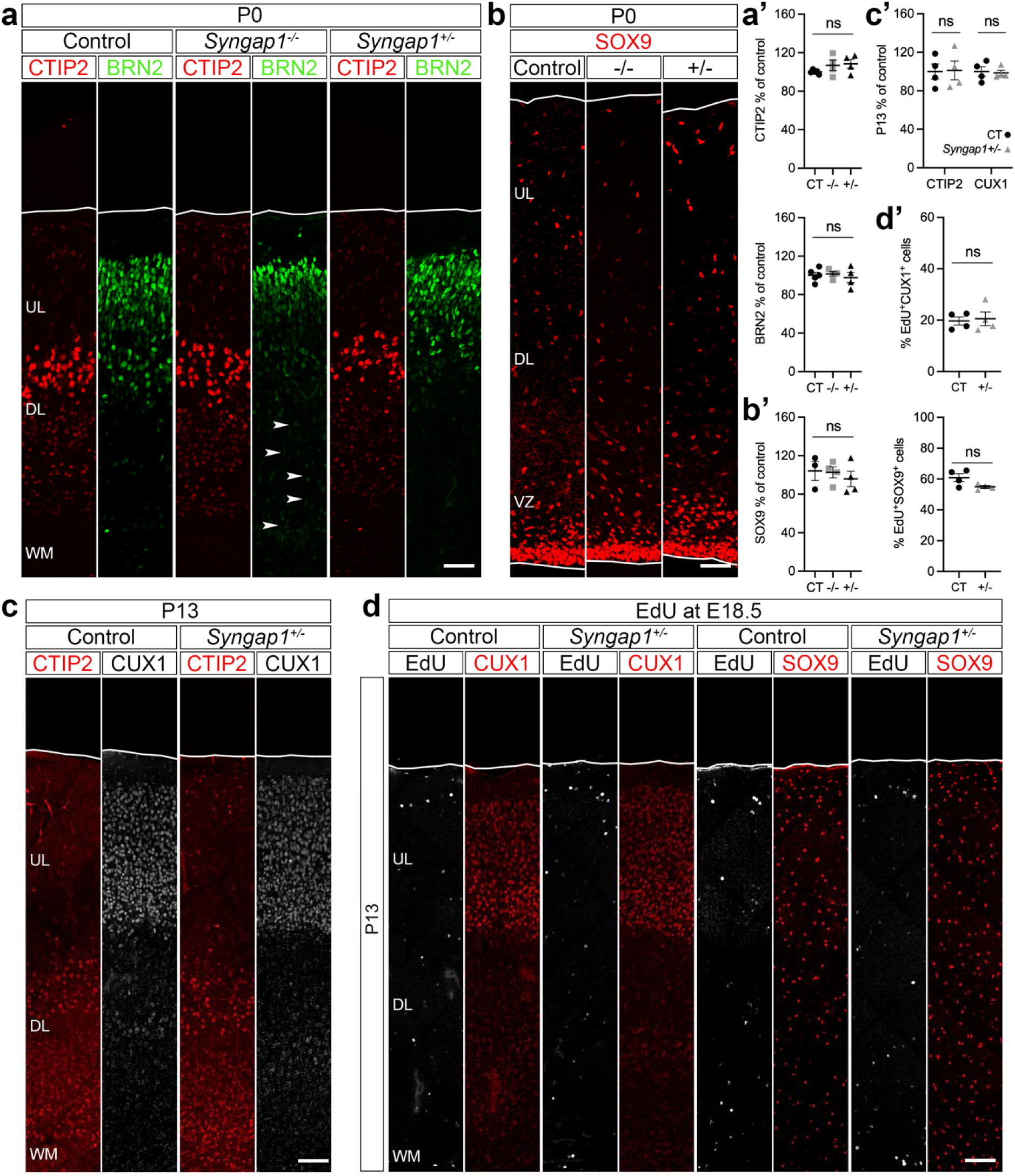
*Syngap1* levels do not affect the number of neurons and glia cells generated in mice. **a, a’,** CTIP2 (red) and BRN2 (green) immunolabeling in the somatosensory cortex of Control (CT), heterozygous (+/-) and homozygous (-/-) *Syngap1KO* mice at P0 (n=5 CT, n=4 -/- and +/- mice; two-sided unpaired t*-*test: CTIP2 - CT vs -/- p=0.1967; CT vs +/- p=0.0870; BRN2 - CT vs -/- p=0.7649; CT vs +/- p=0.6772). Arrowheads highlight migration defects for BRN2- positive cells. **b, b’,** SOX9 (red) immunolabeling in the somatosensory cortex of Control (CT), heterozygous (+/-) and homozygous (-/-) *Syngap1KO* mice at P0 (n=3 CT, n=4 -/- and +/- mice; two-sided unpaired t*-*test: CT vs -/- p=0.8901; CT vs +/- p=0.5368). **c, c’,** CTIP2 (red) and CUX1 (grey) immunolabeling in the somatosensory cortex of Control (CT) and *Syngap1^+/-^* mice at P13 (n=4 mice/genotype; two-sided unpaired t*-*test: CTIP2 - CT vs +/- p=0.9317; CUX1 - CT vs +/- p=0.8127). **d, d’,** EdU (grey) and CUX1 or SOX9 (red) immunolabeling at P13 in the somatosensory cortex of Control (CT) and *Syngap1^+/-^* mice after EdU injection at E18.5 (n=4 mice/genotype; two-sided unpaired t*-*test: EdU^+^CUX1^+^ CT vs +/- p=0.7904; EdU^+^SOX9^+^ CT vs +/- p=0.0690). DL, Deep Layer; UL, Upper Layer; VZ, Ventricular Zone; WM, White Matter. Values are mean ± SEM; ns, not significant. Scale bars: 50 µm (P0), 100 µm (P13).

To further investigate if the late progenitors are generating normal levels of neurons and glia cells, we analyzed cell-fate by injecting EdU at E18.5 and quantifying the number of EdU-positive cells expressing the indicated cellular markers in the somatosensory cortex of control and heterozygous *Syngap1KO* mice at P13. *Syngap1*-deficiency did not affect the ratio of EdU-positive ULNs (EdU^+^CUX1^+^ cells) or EdU-positive glia cells (EdU^+^SOX9^+^ cells) in mice (Fig. 3d,d’).

Overall, these results show that heterozygous and homozygous *Syngap1KO* mice have normal numbers of progenitor cells that proliferate and differentiate to generate appropriate numbers of neurons and glia. Interestingly, the migration of ULNs was slightly affected in both homozygous *Syngap1KO* and homozygous *Syngap1GAP\** mice (arrowheads on Fig. 3a and Extended Data Fig. 2d; Extended Data Fig. 2f and g) suggesting that *SYNGAP1* may play a role in the migration and circuit integration of ULNs and that this effect is GAP activity dependent.

## Discussion

Recent studies have begun to explore the function of *Syngap1* during the early stages of neurodevelopment. Studies using CRISPR-Cas9 unilateral injections in *Xenopus*^20^ showed that *Syngap1* deficiency results in a decrease in brain size and in the area occupied by neural progenitor cells. Willsey et al. observed an increased ratio of neural progenitor cells to neurons suggesting that *Syngap1*-deficient progenitors differentiate less in this species^20^. More recently, Birtele et al.^21^ demonstrated that *SYNGAP1* influences neocortical neurogenesis by accelerating the differentiation of progenitor cells in human cortical organoids. In this context, *SYNGAP1* haploinsufficiency also leads to a reduced ventricular area but the ratio of progenitor cells to neurons is decreased as *Syngap1*-deficient progenitors are more differentiative in human organoids. Our findings present a significant contrast to these studies. While we observed minor ULN migration defects in mice with *Syngap1* haploinsufficiency, we did not observe any alterations in the properties of their neocortical progenitor cells. This discrepancy emphasizes the complexity of *SYNGAP1*’s role in neurodevelopment and suggests a nuanced interaction with species-specific developmental pathways.

The differences in our results may arise from several factors, including variations in experimental models, methodologies, or even the specific developmental stages examined. Willsey et al. using *Xenopus*^20^, while a very valuable model for studying basic aspects of neurodevelopment, limited the study of the more evolved processes of neocortical neurogenesis given the frog has a much simpler brain when compared with mice or humans^42^. Birtele et al. mainly utilized human organoids^21^, which mimic human brain development more closely than mouse models, allowing for insights into the human-specific roles of *SYNGAP1*. Conversely, our study focused on mouse models^8,10^, a traditional but well-characterized system for studying neurodevelopment. This highlights the potential for species-specific mechanisms that govern the biology of progenitor cells and suggests that the role of *SYNGAP1* may differ between species. Although Birtele et al.^21^ described that the ratio of radial glial progenitors to neurons was reduced in *Syngap1KO* mice mimicking the increased differentiating capacity of human radial glial progenitors in *SYNGAP1* haploinsufficient human organoids, the observed differences may have resulted from a comparison between different regions of the rostral-caudal axis in control and *Syngap1KO* brains. Here, we have found that the number of progenitors and the cortical thickness occupied by these progenitors is normal across the rostral-caudal axis in both *Syngap1KO* and *Syngap1GAP\** mice when compared to the corresponding rostral-caudal axis in their control littermates. We showed that the number of self-renewing and differentiative murine progenitors remains normal in *Syngap1KO* at two different developmental ages. Additionally, cortical thickness and the numbers of neurons and glia are not altered in either *Syngap1KO* or *Syngap1GAP\** mice compared to their control littermates. Importantly, despite severe intellectual disabilities, human SYNGAP1 haploinsufficiency patients are rarely diagnosed with structural defects in the brain^17,18^, supporting the generality of our findings.

It is also important to recognize the potential implications of our findings in the broader context of neurodevelopmental disorders associated with *SYNGAP1* mutations. Although research with mice has proven to be instrumental to understand brain function and disease mechanisms, the pathology of neurological disorders is often difficult to fully mimic in mouse models. This divergence is likely due to the evolutionary transcriptomic pathways that separate the early stages of neurodevelopment in rodent’s brain from the human’s brain^43–48^. Supporting this hypothesis, recent evidence shows that certain neurodevelopmental disorders, such as microcephaly, that cannot be mimicked in mice can be replicated in more evolved models like the ferret^49^. Moreover, recent studies have identified human-specific genes and enhancers that differentially affect the biology of progenitor cells and might be associated with the increased neocortical complexity and disease-susceptibility of the human brain^43–48^. Microcephaly-associated genes are included in the list of genes that contain these human-specific enhancers^46^. Like the microcephaly-associated genes, *SYNGAP1*’s involvement in the proliferation and differentiation of progenitors could be more pronounced during critical periods of development in humans, where the demands of cortical assembly and organization are heightened.

Alternatively, *SYNGAP1*’s involvement in the biology of progenitors could be cell specific. Although Birtele et al.’s conclusions mainly refer to the biology of apical progenitors residing in the VZ^21^, *SYNGAP1* might play important roles in outer radial glial cells (oRGCs). ORGCs are neocortical progenitors that reside in the outer SVZ and, although their number and contribution to neurogenesis is very reduced in lissencephalic mice, they are very abundant and account for most neurons in more evolved species with a more complex neocortex^26,32,50–55^. The biology of these cells can be recapitulated in human organoids^56^, and therefore, this might explain some of the differences we observe in mice where oRGCs are present in very limited numbers. This underscores the need for further investigations into the mechanistic pathways influenced by *SYNGAP1* across different species, cell types, and developmental contexts.

In conclusion, our study contributes to the growing body of literature exploring the multifaceted roles of *SYNGAP1* in neurodevelopment. While our results differ from those of Willsey et al.^20^ and Birtele et al.^21^, we emphasize the need for continued research into the diverse mechanisms of *SYNGAP1* across species. Understanding these differences will be critical for developing targeted therapeutic strategies for neurodevelopmental disorders associated with *SYNGAP1* mutations. Collaboration and dialogue within the scientific community will enhance our understanding of these complex processes, and we look forward to further investigations that will bridge the findings from human and animal studies.

## Methods

All the experiments reported in this study adhered to the ethical regulations and guidelines of the National Institute of Health Guide for Care and Use of Laboratory Animals. The experiments involving mice were approved by the Institutional Animal Care and Use Committee at Johns Hopkins University School of Medicine under the study protocol numbers: MO22M217, M023M70, and MO23M52.

### Mice

All animal experiments adhered to the guidelines of the National Institute of Health and were approved by the Institutional Animal Care and Use Committee at Johns Hopkins University School of Medicine. Mice were maintained on a 14h light/10h dark cycle. Both male and female mice were used, and no obvious differences between the sexes were noted. All mice were group-housed in pathogen-free facilities with regulated temperature and humidity and given *ad libitum* access to food and water. All mice used were seemingly free of infection, health abnormalities or immune system deficiencies and were employed independently of their gender. None of the mice used had been used for previous experiments. The date of the vaginal plug detection was designated E0.5, and the date of birth P0. *Syngap1^+/-^* heterozygous knockout mice^10^, *Syngap1* GAP mutants (*+/GAP\**, *GAP*/GAP\**)^8^, and WT (*Control*) littermates were maintained on a mixed background of C57/B6J and 129/SvEv background. *Emx1-Cre* [B6.129S2-*Emx1tm1(cre)Krj*]^57^ and *Brn1/2* conditional knockout mice have been described^40^. CD1 time-pregnant wild type mice were obtained from Charles River Laboratories.

### Animal Euthanasia

We used the most humane methods for euthanasia following the recommendations of the panel report on Euthanasia of the American Veterinary Medical Association and the guidelines of the National Institute of Health Guide for Care and Use of Laboratory Animals. Different methods were used for mice of different ages, according to the experiments that needed to be performed: neonatal and adult mice – isoflurane inhalation followed by cervical dislocation; newborn mice – chilling and decapitation; embryos 14 and 18 days post coitum – chilling and decapitation; newborn to adult mice – cardiac perfusion under full anesthesia.

### RNAscope multiplex *in situ* hybridization

Brain sections were processed for multiplex fluorescent *in situ* hybridization RNAscope^58^ following the manufacturer’s instructions (#323110, Advanced Cell Diagnostics). Gene-specific probes: *Syngap1* (417381), *Pax6* (412821-C2).

### Immunohistochemistry

Embryonic brains were fixed in 4% paraformaldehyde (PFA) in phosphate-buffered saline (PBS) for 2-4h at 4°C. Postnatal mice were transcardially perfused with 20 ml ice-cold 4% PFA using a peristaltic pump at a rate of 2 ml/min. Brains were removed from the skull and postfixed in 4% PFA overnight at 4°C. Embryonic brains were cryopreserved with 30% sucrose, embedded in Tissue-Tek O.C.T. compound (#4583, SAKURA) and frozen at -80°C. They were then sectioned coronally at 16 µm with a cryostat (CM3050 S, Leica). Postnatal brains were embedded in 3% low melting point agarose (#R0801, Thermo Fisher Scientific) in PBS and sectioned coronally at 60 µm with a vibrating microtome (VT1200S, Leica). For immunohistochemistry, brain sections were permeabilized in PBS containing 0.2% Triton X-100 and blocked in 10% goat serum for 1h at room temperature. Brain slices were then incubated with primary antibodies in blocking solution overnight (or 48h depending on the thickness of the section and antibody used) at 4°C and subsequently incubated with the appropriate secondary antibodies diluted in blocking solution for 2h at room temperature. To label all cell nuclei, the fluorescent nuclear dye DAPI (1 μg/ml, #D9542, Sigma-Aldrich) was included with the secondary antibody solution. The sections were mounted with ProLong^TM^ Gold (#P36930, Thermo Fisher Scientific). Primary antibodies: anti-CTIP2 (1:1000, rat monoclonal, ab18465 Abcam); anti-CUX1 (1:1000, rabbit polyclonal, 11733-1-AP Proteintech); anti-BRN2 (1:1000, rabbit monoclonal, 12137 Cell Signaling); anti-Ki67 (1:500, rabbit polyclonal, ab15580 Abcam or 1:500, rat monoclonal, 14-5698-82 Thermo Fisher Scientific); anti-PAX6 (1:250, mouse monoclonal, MA1-109 Thermo Fisher Scientific or 1/500, rabbit polyclonal, 901301 Biolegend); anti-TBR2 (1:500, rat monoclonal, 14-4875-82 Thermo Fisher Scientific or 1:500, rabbit polyclonal, ab183991 Abcam); anti-SOX9 (1:1000, rabbit monoclonal, ab185966 Abcam); anti-PH3 (1:1000, rat monoclonal, ab10543 Abcam); anti-γ-TUBULIN (1:500, goat polyclonal, A. Holland gift^59^); anti-Centrin (1:500, rabbit polyclonal, A. Holland gift^60^). Secondary antibodies: Alexa 555, 647, 488 anti-rat (A21434; A21247, A11006), anti-rabbit (A21430; A21246) and anti-mouse (A21425; A21237) and Alexa 488 anti-rabbit (A11070); all diluted 1:500 and all from Thermo Fisher Scientific. Antigen retrieval^61^ was performed for all immunohistochemistry at embryonic ages. The method used here was a heat-induced citrate method^61^. The sections were then mounted with ProLong^TM^ Gold (P36930, Thermo Fisher Scientific).

### EdU injections and labeling

EdU (5-ethynyl-2’-deoxyuridine; #A10044, Thermo Fisher Scientific) was diluted at 10mg/ml in 0.9% NaCl (#114-055-101, Quality Biological) and administered in pregnant females at the specified embryonic pregnancy stage at 50mg/Kg body weight. For proliferation analysis, a single intraperitoneal EdU injection was administered at E18.5, embryos were fixed after 1h and the number of EdU^+^ cells was quantified. For cell cycle exit calculation, a single intraperitoneal EdU injection was administered at E17.5, embryos were fixed after 24h and the percentage of EdU^+^ cells labeled with Ki67 was calculated. For cell-fate analysis, a single intraperitoneal EdU injection was administered at E18.5, and the number of EdU^+^ cells expressing the indicated cellular markers were analyzed at P13. EdU was labeled using Click-iT^®^ EdU imaging kits with alexa 555 (#C10638, Thermo Fisher Scientific). Briefly, brain sections were post-fixed with 3.7% formaldehyde and permeabilized with 0.5% Triton X-100 in PBS. The Click-iT^®^ reaction cocktail was then added, and the sections incubated in the dark for 30 minutes at room temperature. After the washes, the brain sections were stained for the different proteins of interest using the immunohistochemistry protocol previously described.

### Cleavage plane analysis

PH3 immunostaining was used to identify mitotic cells. γ-TUBULIN immunostaining was used to determine the position of both centrosomes (γ-TUBULIN positive) in mitotic cells and to mark the cleavage plane in relation to the ventricular surface. The angle of the cleavage plane was measured using ImageJ.

### Imaging

All immunohistochemistry and ISH RNAscope data were acquired in an equivalent latero-medial level of the cortex from at least 3 independent animals per condition or genotype. Images were captured using a Zeiss LSM 800 confocal microscope. Maximum intensity projections were generated in ImageJ or Imaris software. In all fluorescence microscopy figures, different channels of image series were represented in pseudocolor, and contrast and brightness were adjusted manually using ImageJ or Adobe Photoshop software. All markers were counterstained with DAPI to allow visualization of overall cellular density.

### Histological analysis

A series of *z*-stack (depth of 10 μm) confocal tiled images were used for cell quantification; 150 μm-wide images (embryonic brains) or 300 μm-wide images (postnatal brains) comprising the entire extension of the cortex were analyzed. All quantification were made in an equivalent latero-medial level of the cortex from at least 3 independent animals per condition or genotype. The number of cells expressing the markers of interest was quantified in radial sections of the same latero-medial level and adjusted per area. Cortical thickness was analyzed by measuring the length from the lower limit of the VZ to the upper limit of the CP. PAX6^+^ or TBR2^+^ cortical thickness was analyzed by measuring the length occupied by PAX6^+^ or TBR2^+^ cells from the lower limit of the VZ into the CP. To analyze the distribution of BRN2^+^ cells we divided the cortex into longitudinal bins, a rectangle was drawn from the border of the white matter with layer VI to the pial surface. This rectangle was then divided into ten equal bins and the number of cells within each bin was quantified.

### Statistical analysis

Statistical analyses of the immunohistochemistry were done using GraphPad Prism software. Statistically significant differences were assessed by Student’s unpaired t-test. To compare the distribution of BRN2^+^ cells in the different bins of the cortex for the different genotypes we used two-way analysis of variance (two-way ANOVA-Šídák’s multiple comparisons test; F_DFn, DFd_). P<0.05 was considered a significant difference. All values represent individual animals mean ± standard error of the mean (SEM). The statistical test used, and the statistical significance are indicated in figure legends.

## Acknowledgments

We thank members of the Huganir and Müller laboratories, including Yoichi Araki, for suggestions, reagents and technical help; Andrew Holland for the g-TUBULIN and Centrin antibodies. We are grateful to Ashley Irving, Sarah Rodriguez, Michelle Monroe, Tajma Smith and Trinity Walker for mouse maintenance and genotyping assistance. We thank Giorgia Quadrato for her suggestions and constructive discussions.

## Funding

This work was supported by National Institutes of Health grants R37NS036715 (RLH), R01HG012357 and RF1MH121539 (UM).

## Data and materials availability

All data are available in the main text or the Extended Data. Correspondence and requests for materials should be addressed to RLH.

## Author Contributions Statement

Conceptualization: SB, IH, UM and RLH; Methodology: SB, IH, UM and RLH; Investigation: SB, IH, UM and RLH; Validation: SB, IH and RLH; Visualization: SB and IH; Formal analysis: SB, IH, UM and RLH; Data curation: SB and IH; Resources: UM and RLH; Supervision: UM and RLH; Project administration: UM and RLH; Funding acquisition: UM and RLH; Writing: SB, IH, UM and RLH.

## Competing Interests Statement

The authors declare no competing or financial interests.

## Extended Data

**Figure 1.**
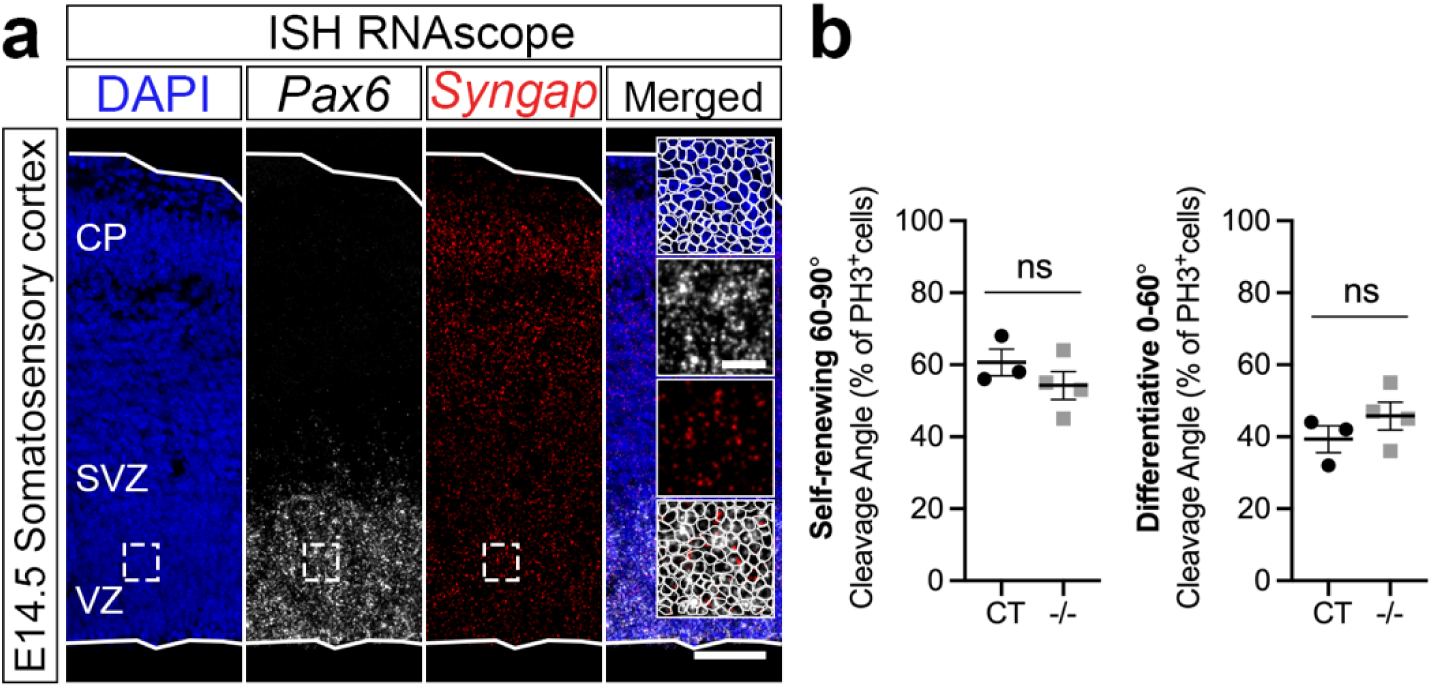
*Syngap1* expression and progenitors’ cleavage plane analysis at E14.5. **a,** RNAscope for *Syngap1* (red) and *Pax6* (grey) in the somatosensory cortex of wild-type mice at E14.5. Low and top lines represent the limits of the ventricular zone (VZ) and cortical plate (CP), respectively. Boxed area at higher magnification on the right. Lines circulating the cells show co-expression *Syngap1* and *Pax6* in the VZ. **b,** Cleavage plane analysis in Control (CT) and homozygous (-/-) *Syngap1KO* ventricular zone (VZ) at E14.5 (n=3 CT, n=4 -/- mice; two-sided unpaired t*-*test: CT vs -/- p=0.3010). SVZ, Subventricular Zone. Values are mean ± SEM; ns, not significant; Scale bars: 50 µm (lower magnification), 10 µm (higher magnification).

**Figure 2.**
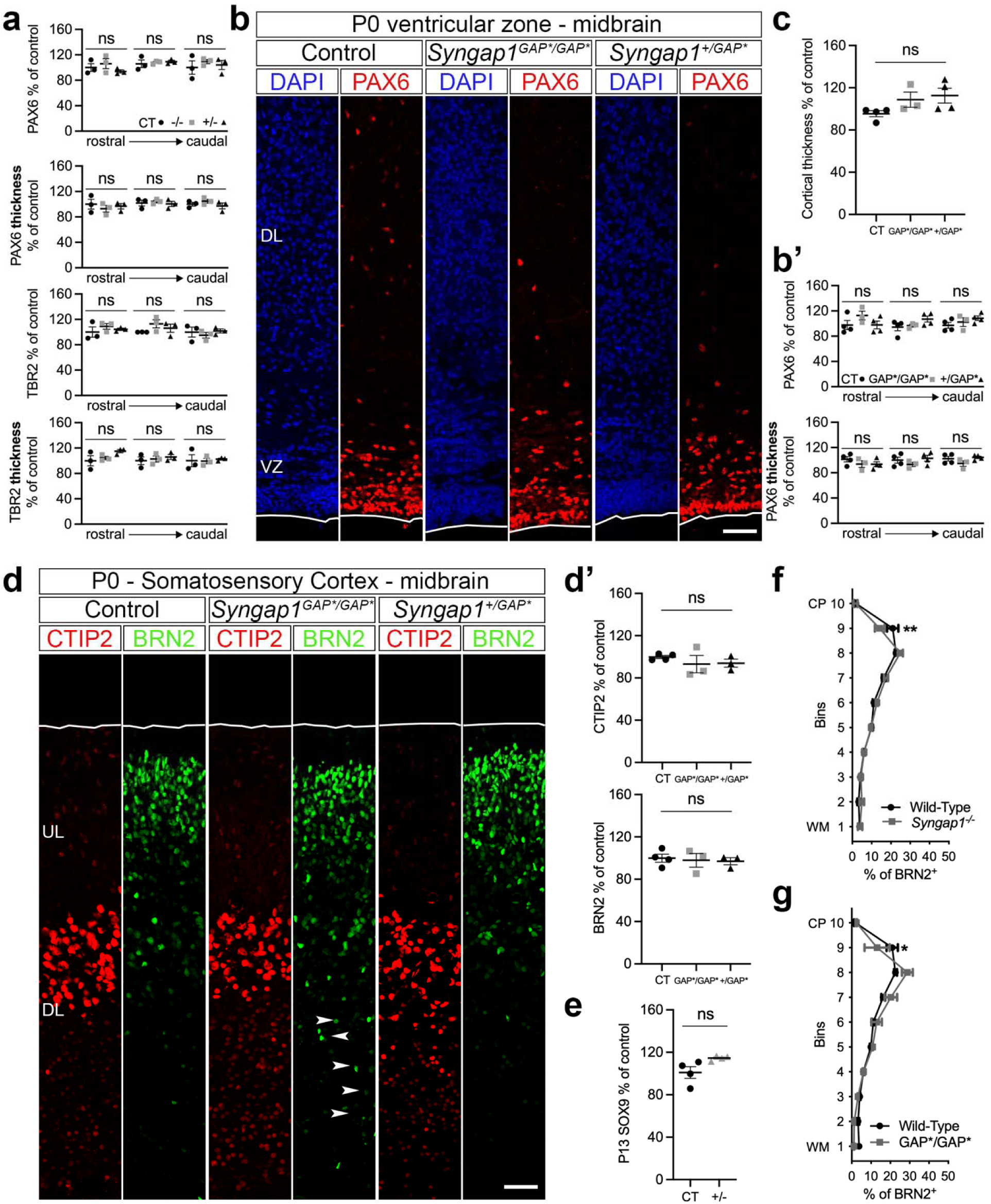
*Syngap1* levels or activity do not affect the number, properties or output of progenitor cells in mice. **a,** PAX6^+^ and TBR2^+^ cells quantification along the rostral-caudal axis in Control (CT), heterozygous (+/-) and homozygous (-/-) *Syngap1KO* somatosensory cortex at E18.5 (related to Fig. 1c; n=3 mice/genotype; two-sided unpaired t*-*test: PAX6^+^ cells: rostral – CT vs -/- p=0.5678; CT vs +/- p=0.4186; middle – CT vs -/- p=0.7337 CT vs +/- p=0.6224; caudal – CT vs -/- p=0.4413; CT vs +/- p=0.7667; PAX6^+^ cortical thickness: rostral – CT vs -/- p=0.4893; CT vs +/- p=0.7305; middle – CT vs -/- p=0.6480; CT vs +/- p=0.8375; caudal – CT vs -/- p=0.3311; CT vs +/- p=0.6496; TBR2^+^ cells: rostral – CT vs -/- p=0.3851; CT vs +/- p=0.6316; middle – CT vs -/- p=0.1170; CT vs +/- p=0.4029; caudal – CT vs -/- p=0.6087; CT vs +/- p=0.8486; TBR2^+^ cortical thickness: rostral – CT vs -/- p=0.6297; CT vs +/- p=0.1452; middle – CT vs -/- p=0.7394; CT vs +/- p=0.4585; caudal – CT vs -/- p=0.9664; CT vs +/- p=0.7995). **b,** PAX6 (red) immunolabeling in the somatosensory cortex of Control (CT), heterozygous (+/*GAP\**) and homozygous (*GAP*/GAP\**) mice for *GAP-AL* mutation (*Syngap1GAP\**) at P0 (n=4 CT and +/*GAP\**, n=3 *GAP*/GAP\** mice; two-sided unpaired t*-*test: CT vs *GAP*/GAP\** p=0.7715; CT vs +/*GAP\** p=0.1392). **b’,** PAX6^+^ cells quantification along the rostral-caudal axis in the somatosensory cortex of Control (CT), heterozygous (+/*GAP\**) and homozygous (*GAP*/GAP\**) mice for *GAP-AL* mutation (*Syngap1GAP\**) at P0 (n=4 CT and +/*GAP\**, n=3 *GAP*/GAP\** mice; two-sided unpaired t*-*test: PAX6^+^ cells: rostral – CT vs *GAP*/GAP\** p= 0.2071; CT vs +/*GAP\** p=0.9970; middle – CT vs *GAP*/GAP\** p= 0.7530; CT vs +/*GAP\** p=0.1392; caudal – CT vs *GAP*/GAP\** p= 0.5445; CT vs +/*GAP\** p=0.1065; PAX6^+^ cortical thickness: rostral – CT vs *GAP*/GAP\** p= 0.3185; CT vs +/*GAP\** p=0.2230; middle – CT vs *GAP*/GAP\** p= 0.3356; CT vs +/*GAP\** p=0.6625; caudal – CT vs *GAP*/GAP\** p= 0.2457; CT vs +/*GAP\** p=0.6263). **c,** Cortical thickness in the somatosensory cortex of Control (CT), heterozygous (+/*GAP\**) and homozygous mice for *GAP-AL* mutation (*Syngap1GAP\**) at P0 (n=4 CT and +/*GAP\**, n=3 *GAP*/GAP\** mice; two-sided unpaired t*-*test: T vs *GAP*/GAP\** p= 0.1116; CT vs +/*GAP\** p=0.0644). **d, d’,** CTIP2 (red) and BRN2 (green) immunolabeling in the somatosensory cortex of Control (CT), heterozygous (+/*GAP\**) and homozygous (*GAP*/GAP\**) mice at P0 (n=4 CT, n=3 *GAP*/GAP\** and +/*GAP\** mice; two-sided unpaired t*-* test: CTIP2 - CT vs *GAP*/GAP\** p=0.3848; CT vs +/*GAP\** p=0.1743; BRN2 - CT vs *GAP*/GAP\** p=0.7894; CT vs +/*GAP\** p=0.6107). Arrowheads highlight migration defects for BRN2-positive cells. **e,** SOX9 quantification in the somatosensory cortex of Control (CT) and *Syngap1^+/-^*mice at P13 (n=4 mice/genotype; two-sided unpaired t*-*test: CT vs +/- p=0.0528). **f, g,** Distribution of BRN2^+^ cells in the somatosensory cortex of Control (CT), *Syngap1^-/-^* (**f**; related to Fig. 3a) and *Syngap1^GAP*/GAP*^* (**g**; related to Extended Data Fig. 2d) mice at P0 (n=4 CT and *Syngap1^-/-^*, n=3 *Syngap1^GAP*/GAP*^*mice; two-way ANOVA-Šídák’s multiple comparisons test: *Syngap1^-/-^*– 9-p=0.0017, F_9,60_=2.236; *Syngap1^GAP*/GAP*^* – 9-p=0.0415, F_9,50_=2.116). DL, Deep Layer; UL, Upper Layer; VZ, Ventricular Zone; WM, White Matter. Values are mean ± SEM; ns, not significant. Scale bars: 50 µm (P0), 100 µm (P13).

## Notes

### Competing Interest Statement

The authors have declared no competing interest.

